# Immediate early gene kakusei plays a role in the daily foraging and learning of honey bees

**DOI:** 10.1101/760587

**Authors:** Asem Surindro Singh, Pamela Cappelletti, Marco Feligioni

## Abstract

*kakusei* is a non-coding RNA that is overexpressed in foraging bee brain. This study describes a fundamental role of the IEG *kakusei* during the daily foraging of honey bees. *kakusei* was found to be transiently upregulated within two hours during rewarded foraging. Interestingly, during unrewarded foraging the gene was also found to be up-regulated, but immediately lowered as food was not rewarded. Moreover, the *kakusei* overexpression was diminished within a very short time when the time schedule of feeding was changed. This indicates the potential role of *kakusei* on the motivation of learned reward foraging, reflecting associative learning memory. These results provide evidence for a dynamic role of *kakusei* during foraging of bees, and eventually its possible involvement in learning and memory. Thus the *kakusei* gene could be used as key search tool in finding distinct molecular pathways that mediate diverse behavioral components of foraging.

## Introduction

The systematically well organized social behavior of honey bee foraging has been an attractive field of research. Further exploration of the dynamics of honey bee foraging at the molecular and cellular level could help in uncovering the complex mechanisms of social behavior. Honeybee foraging comprises several behavioral components including learning, memory, social interaction and communication. In 1973 the Austrian ethologist Karl Ritter Von Frisch (Austrian ethologist) was honored with Nobel Prize in Physiology or Medicine for his investigations of sensory perceptions in honey bees [1]. He translated the meaning of the bee waggle dance which represents a particular movement of honey bee that looks like the figure of numeric eight, and revealed that with the help of this dance foraging bees of the same colony share information, including about the direction and distance to patches of flowers yielding nectar and pollen, sources of water, and new nest site locations. [2-4]. Bee foraging has gained enormous attention, since it offers the opportunity to elucidate the complexity of behavioral mechanisms involved in accomplishing the foraging task. Thanks to the efforts of several researchers, today we have a substantial amount of knowledge on this subject. However, understanding of the context of molecular and cellular mechanisms underlying foraging tasks is still poor.

During foraging, bees fly back and forth several times during the day between the hive and the food location, collecting nectar/pollen and bringing it to the hive for their colony. Foraging involves highly systematic and dynamic behavioral capacities that include long distance navigation using the sun as compass, evaluation of food quality, learning and memorizing flower cues, sense of colour, social communication/interaction for coordination in collecting nectar, pollen, and water. [2,5,6].

For decades the outdoor experiments have been routinely performed by feeding bees on pollen or sucrose solution placed in a specific place in order to mimic the natural foraging environment in close proximity. Using this experimental setup, we attempted to identify foraging regulatory genes in the European honey bee species *Apis mellifera* while the bees were performing their daily routine foraging. For this, the immediate early genes (IEGs) were the ultimate targets to examine, as they are well known as neural markers.

IEGs are rapidly induced by a large number of stimuli, and alterations of their expression are considered as part of the general neuronal response to natural stimuli as interpreted by normal synaptic activity [7]. The products of IEGs can activate downstream targets that typically function as part of a network of constitutively expressed proteins [7]. IEGs are also known to be the first activated genes that link cellular membrane events to the nucleus [8], and the gene expression changes are required for the late neuronal response which relates to in the process of learning and memory formation [9] and which is a part of everyday brain functions [10]. In addition, depending on the type of the stimuli, the IEG-encoded proteins can be individually regulated in different parts of the brain [8,11], indicating that the same or different IEGs, when expressed in different parts of the brain, regulate different functions. In our recent work, we have demonstrated involvement of two IEGs i.e., early growth response 1 (*egr-1*), nuclear hormone receptor 38 (*hr38*) and their corresponding partner genes such as ecdysone receptor (*ecr*), dopamine/ecdysteroid receptor (*dopecr*), dopamine decarboxylase (*ddc*) and dopamine receptor 2 (*dopr2*) during the daily foraging of bees [12]. Other papers reported on the involvement of *egr-1* in the transition from nursing to foraging bees [13] and the role of *hr38* in caste and labor division [14], while the other genes were shown to be act as *egr-1* downstream genes involved in ecdysteroid and dopamine signaling with a role in the processing of courtship memory [15] in Drosophila and in olfactory learning and memory [16] in honey bees, respectively. The IEG *c-jun* (also known as jun-related antigen, *jra* in fruit fly) and *egr-1* have been used as neuronal markers for the identification in honey bees of specific brain regions involved in the time memory as well as in, innate and learned behaviours [17,18,19]. Interestingly, while most of the above mentioned genes are already well known through various research studies, a recently discovered insect IEG *kakusei*, a noncoding RNA, has been found to have function in the patterning of neuronal activity in the foraging bees [20,21,22]. Expression of *kakusei* was detected during ice-induced seizures in bees awakening from anesthesia, and its expression was prominent in the Kenyon cells of the mushroom bodies [20,21,22]. Since *kakusei* was found as neural marker, we were interested to extend the work on this gene in our experimental design.

The present study represents further investigation of immediate early genes (IEG) that could play a role during the daily foraging of honey bees, and is an expansion of our previous report by Singh et al [12]. We selected two potential IEGs *kakusei* and *c-jun*, and four other orthologous genes that have been reported to be involved in cognitive process in vertebrates: extracellular signal-related kinase (*rk7*), glutamate receptor (*GluR*), 5-hydroxy tryptamine (serotonin) receptor 2 alpha (*5-HT2α*) and dopamine receptor 1 (*Dop1*). Among the six genes tested, we observed only for the kakusei gene having transient and prolonged upregulation during reward foraging and a short period of overexpression during unrewarded foraging. These findings demonstrate a potential role for *kakusei* during foraging of bees, and that *kakusei* may be important for associative learning while foraging.

## Experimental procedures

### Foraging experiment and sample collection

The behavioral tests were performed by using an outdoor flight cage (length=12m, height=5m, width=2.5m) located within the NCBS campus, Bangalore, India. The *Apis mellifera* colonies were kept within the outdoor flight case and bees were fed with pollen and 1 M sugar solution placed on a green and yellow plastic plate respectively. The distance of feeders was 10m from the beehive and with 1.5m gap between the two feeders. The feeding time was from 14.00 to 17.00 hrs every day. Sample collection was started after the foraging bees had visited feeders for several days. For the gene expression profiling, nectar foragers were collected and the gene expression analysis was carried out with those samples. The procedures were previously described [12].

### Collection procedure and sample grouping

Collection during foraging: The first arriving foragers were caught at the feeder plate before presenting the sugar solution on the feeder (0min group) and the caught bees were immediately flash frozen in liquid nitrogen for further gene expression analysis. Soon after the first collection, sugar solution was presented and the first arriving foragers were marked by paint markers. The marked bees were collected at a series of time points with 15 min intervals up to 2 hrs (15min, 30min, 45min, 60min, 75min, 90min, 105min, and 120min groups), then flash frozen immediately in liquid nitrogen. We caught paint-marked foragers as soon as they landed over the plate and before they start drinking the sugar solution at those set times during their repeated trips. In one day we collected about 24 bees, 1-2 bees for each time point starting from 14.00 hours until 16.00 hours and continued in following days until we obtained 5 bees at each time point.

Collection before and after foraging: Pre-foraging bees were paint-marked while foraging and collected in the morning (09.00 hrs) of the next day in the hive before they started foraging; whereas post-foraging bees were caught in the hive in the evening (09.00 hrs later) after the bees finished foraging, on the same day the bees were paint marked. We took gentle care during the collection procedure in order that the foraging bees were not disturbed and to avoid inducing stress phenomena; thereby the interaction between the collector and the bees were considered minimal [23,24,25]. The caught bees were immediately flash frozen in liquid nitrogen.

Collection without food reward: This category included only the foraging bees collected at the empty feeder plate at different time points. The foragers were marked at 0min upon their arrival at the empty feeder and 0min samples were collected. After the collection of 0min samples, 1 M sucrose solution was presented to let the bees continue coming and the collection continued for one hour with samples at 15min, 30min, 45min and 60min. While the unmarked bees were drinking, the marked bees were caught as soon as they landed on the feeder plate; it was made sure that those bees had not touched the sucrose solution. About 2 bees were collected in each day of collection and the collection started at 14.00 hrs. The bees were immediately flash frozen in liquid nitrogen as soon as they were caught.

Collection at different feeding time: Here the procedure is same with that of (1). The only difference was that the feeder was presented at 11.00 hrs instead of the normal feeding time 14.00 hours, and the collection also started from the same time and continued for 1 hr with samples at 0min, 15min, 30min, 45min and 60min.

### Brain dissection, RNA and cDNA preparation, and qPCR

The frozen bees were lyophilized at -500C with vacuum at 0.420 mBar for 20min, using a Freeze Zone® PlusTM 4.5 liter cascade Freeze Dry System (Labconco Corporation, Kanas City). Brain dissection was performed in a glass chamber containing 100% ethanol placed on a dry ice platform. The whole brain from each bee was dissected and placed into a micro centrifuge tube separately, and 500 µl Trizol (Trizol® Reagent, ambion RNA, life technology) was added. The brain was homogenized, RNA was extracted, and cDNA was prepared from that RNA, using the kits supplied by Invitrogen (Thermo Fisher Scientific) following the manufacturer’s protocols. The cDNA from each brain was subjected to qPCR using a 7900HT Fast Real Time PCR System (Applied Biosystem, Singapore) in a 10µl reaction volume containing oligonucleotide primers (Sigma Aldrich) specific to target genes and SYBR Green (KAPA Syber® FAST PCR Master Mix (2X) ABI Prism®). The qPCR cycles followed Applied Biosystem protocol and the Rp49 gene was used as endogenous control [26] in each qPCR run and analysis. The details of the target genes and the oligonucleotide primers are provided in Table 1.

### Statistical analysis of qPCR

We calculated relative gene expression level using the relative standard curve method with the help of SDS 2.4 software provided with the 7900HT Fast Real system. The standard deviation was calculated following Applied Biosystem’s ‘Guide to performing relative quantification of gene expression using real-time quantitative PCR’. The fold change was determined relative to time 0, and the statistical significance was tested using one-way ANOVA with Turkey-Kramer post-hoc multiple comparison test; the analysis was carried out with the help of GraphPad InStat software [27]. Normal distribution of each group compared was tested using the D’Agostino & Pearson omnibus normality test. In order to check the differences among the independent experiments, two-way ANOVA program of GraphPad Prism was also employed (GraphPad Software Inc. www.graphpad.com).

## Results

In this study, only the immediate early gene *kakusei* showed a remarkable transient upregulation during the course of food reward foraging. The other five genes *erk7, c-jun, glur, 5-ht2α, dop1* showed no statistically significant differences (Supplementary Figure S1).

### Expression profile of *kakusei* during food reward foraging

The expression of *kakusei* was measured at eleven time points, BF (pre/before foraging), 0min, 15min, 30min, 45min, 60min, 75min, 90min, 105min, 120min and AF (post/after foraging). In order to check the consistency of the results, three experiments were performed using independent samples collected from two different bee colonies over three months. Each of the three experiments demonstrated transient increased of *kakusei* level during the continuous food reward foraging over the collection time of two hours. The results are summarized in Figure 1 (and Supplementary Table S1).

**Figure 1.**
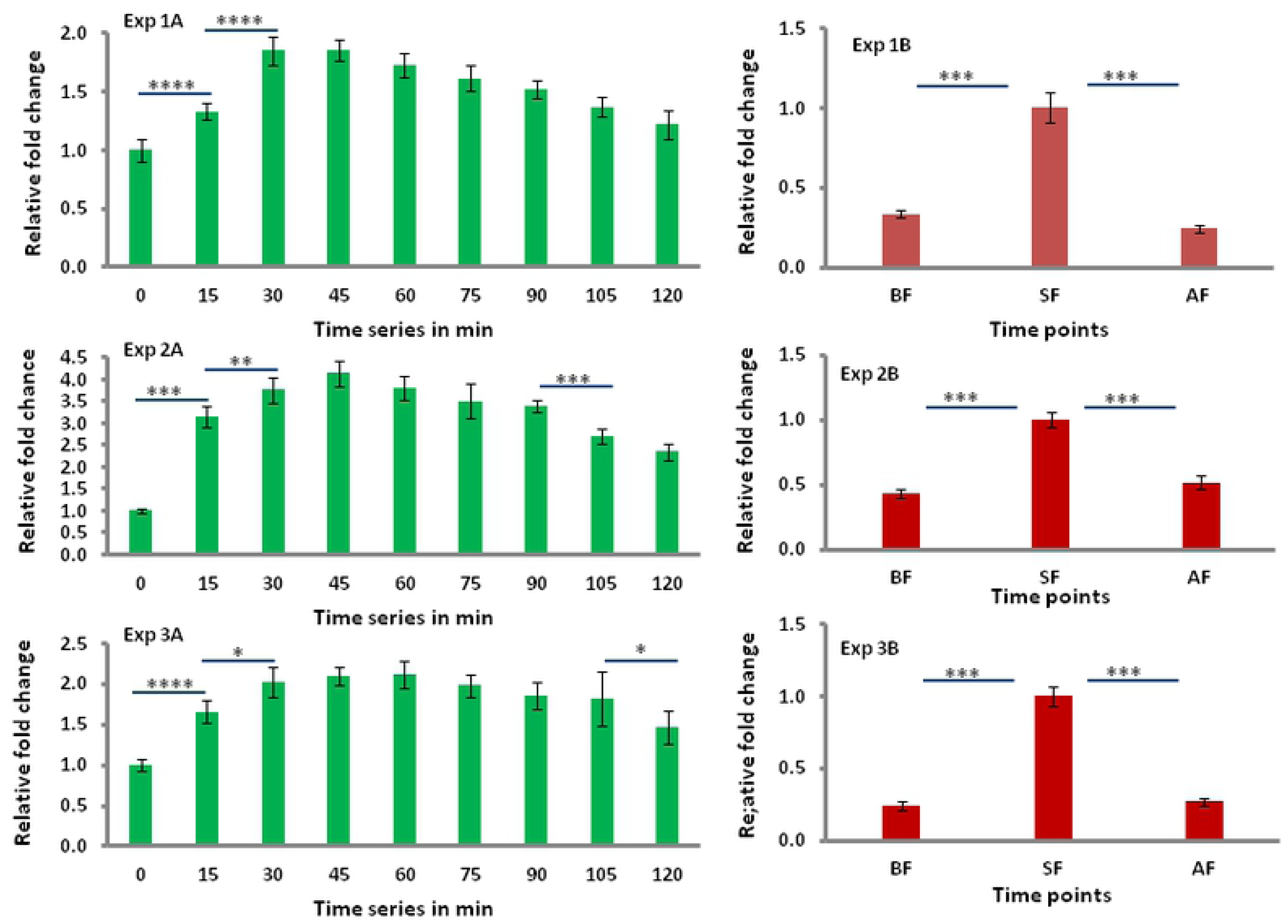
Expression changes of IEG *Kakusie* during daily foraging of bees. Data are shown as fold changes with respect to t0 (mean value was set as 1 at this time point) and each green/red bar graph represents *kakusei* level at each time point. The green bar graphs show the *kakusei* expression level from t0 to t120 whereas red bar graphs represent *kakusei* level difference between the start of foraging (SF/t0: 14.00 hrs) and before-foraging (BF: 09.00 hrs) or after foraging (AF: 18.00 hrs). Each time point has sample size n=5. For statistics one-way ANOVA with Turkey-Kramer post-hog multiple comparison test were performed and number of asterisk symbol represents * P < 0.05, ** P < 0.01, *** P < 0.001, and **** P < 0.0001.

We observed an increase of *kakusei* expression from 0min (start of foraging, 14.00 hrs) to 15min (P = <0.0001) and from 15min to 30min (P = <0.0001) (Figure 1 Exp 1A). While the increase held at about 30min and a decrease was apparent from 45min, there was sustained over expression of *kakusei* further down along the time series until 120min during the reward foraging as compared to the level at 0min (Figure 1 Exp 1A and Supplementary Table S1). The similar pattern was also observed in the subsequent two experiments with independent samples (Figure 1 Exp 2A/3A and Supplementary Table S1). The *kakusei* level was also significantly higher (P = <0.0001) at the start of foraging (0min) when compared to the level at pre foraging (09.00 hrs) or post foraging (18.00 hrs), while there was no significant difference between the before and after foraging samples (Figure 1 Exp 1B). This observation was further validated by the following two independent experiments (Figure 1 Exp 2B/3B). We further examined whether *kakusei* overexpression pattern was statistically different among the three independent results. We found significantly different overexpression level of *kakusei* among the three independent experiments (P=0.0001). On the other hand, the difference of *kakusei* level between Exp-1/Exp-3 vs Exp-2 (P=0.0001) was much higher than Exp-1 vs Exp-3 (P=0.044). In our belief, this difference might be due age difference among the group of bees in Exp-1A/1B, Exp-2A/2B and Exp-3A/3B as we did not maintain the age of foraging bees during sample collection. The graphical results of the analysis are presented in Figure 2A. Subsequently we observed similar result between start of the foraging vs pre/post foraging groups as well, but no statistical difference between pre vs post foraging (Figure 2B).

**Figure 2.**
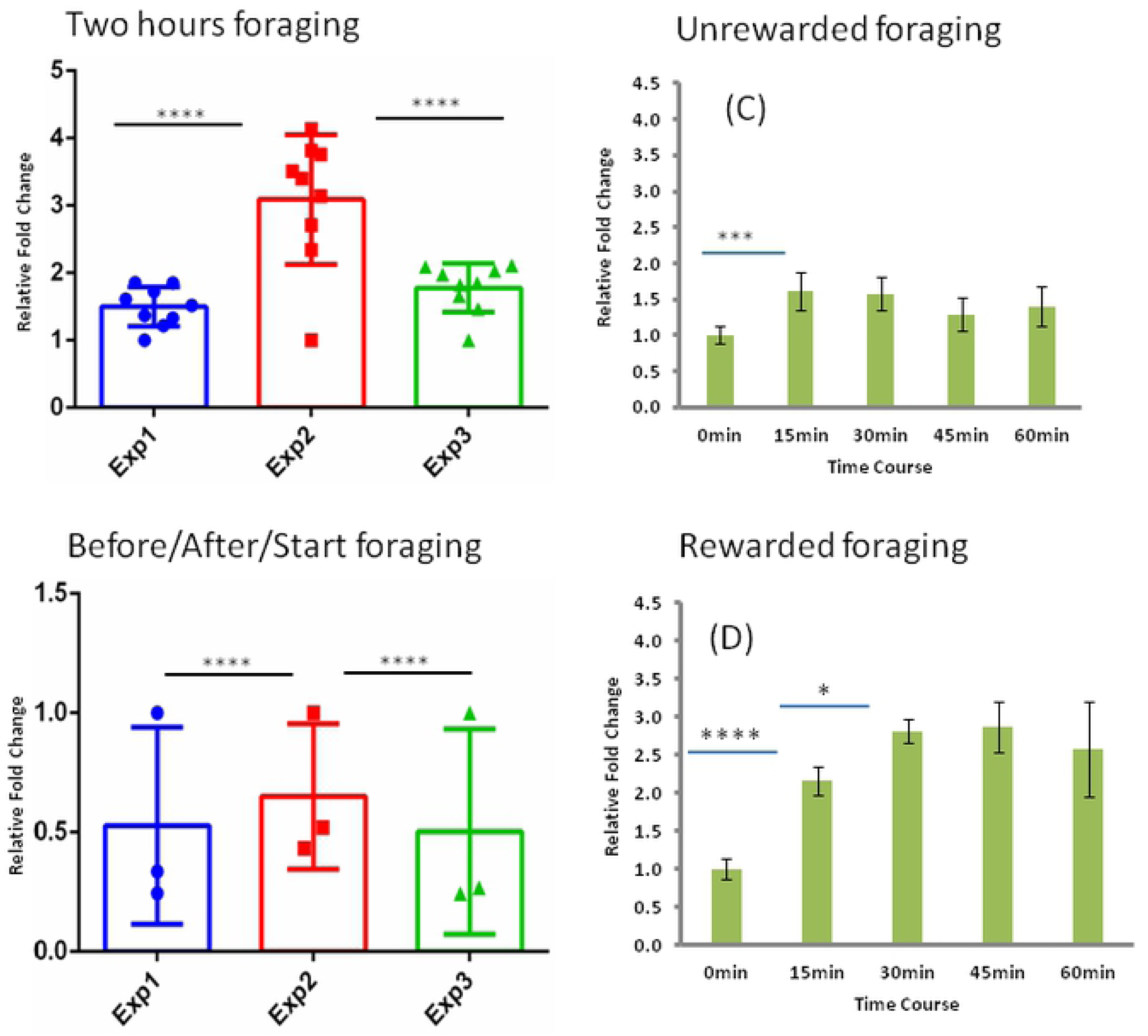
Comparison of three independent experiments of foraging in two hours (A) and before/after foraging (B), and expression changes of IEG *kakusie* with unrewarded (C) and different foragingtime (D). **A&B**: The three independent experiments shown in Figure 1 are examined for interaction using two-way ANOVA. The bars represent mean *kakusei* expression in each experiment and the dots represent the mean level of *kakusei* at each time points. Different colors of bars/dots and different shapes of dots represent for different experiments. **C&D**: The *kakusei* expression profile is presented as fold changes with respect to t0 (mean value was set as 1 at this time point) and each green bar graph represents *kakusei* level at each time point. Figure 2A and 2B represent data for unrewarded and different feeding time foraging respectively. Each time point has sample size n=5. For statistics one-way ANOVA with Turkey-Kramer post-hog multiple comparison test were performed. The number of asterisk symbol represents * P < 0.05, ** P < 0.01, *** P < 0.001, and **** P < 0.0001.

### Expression changes of *kakusei* during unrewarded foraging

In order to confirm whether the *kakusei* upregulation was solely due to the food reward, we presented the empty plate at the usual feeding time 14.00 hrs and the bees were collected without feeding following the procedure described in the methods section. Interestingly, *kakusei* expression from the whole bee brains showed upregulation at 15min (P = 0.0011; Fig 2C), indicating a potential effect on *kakusei* of the learned motivation of food reward. However, the value no longer increased after 15min and the level dropped by 45min (Figure 2C and Supplementary Table S2) indicating that a food reward is required in order to sustain *kakusei* upregulation. This result demonstrates that *Kakuei* is involved in associative learning.

### Testing for time-dependent food reward foraging effect on *kakusei* expression

To further examine whether the *kakusei* expression depends on the time of feeding, the sucrose solution was presented at 11.00 hrs, three hours ahead of the usual feeding time instead of 14.00 hrs (it may be recalled that the bees were fed every day at 14.00 hrs in the above experiments) and the marked bees were collected over the period of 1 hour, at 0min, 15min, 30min, 45min and 60min, as described in method section. A similar pattern of overexpression of *kakusei* was observed as in the previous experiment of food reward foraging in which the collection began at 14:00 hrs. The increase continued from 0min to 15min (P = <0.0001) and 15 min to 30min (P = 0.0195), then no further increase (Figure 2D and Supplementary Table 2). This further reveals that *kakusei* role is independent of feeding time during the day, but responds to the reward of food during foraging.

## DISCUSSION

This study extends our recent report [12] on the search for immediate early genes (IEGs) that could be used as search tool towards finding the molecular and cellular mechanisms underlying social behavior using foraging of honey bees as model system. As the foraging of bees consists of systematically well organized social behaviors that include learning, memory, social interaction and communication etc; honey bee foraging has been extensively studied in various research objectives in connection to social behavior. Noting that honey bees learn food sources, identify the food location, memorize the place, interact and communicate among the foragers, that can be clearly observed during the time of their daily foraging, it is a thoughtful approach to find the genes that initiate and continue foraging and examine their roles in these behavioral routines. Our previous report by Singh et al., 2018 [12], demonstrates that the two IEGs *egr-1, hr38* and their downstream genes such as *ecr, dopecr, ddc*, and *dopr2* are involved in the daily foraging. Moreover, *egr-1* and *hr38* are also involved in associative learning that occurs during foraging. In the present study the role of the newly discovered insect IEG *kakusei* has been described, unraveling its role during the foraging of bees and in associative learning during foraging.

*kakusei* is an insect immediate early gene recently identified by Kubo and his group in the seizure induced in bees by cold or CO_2_ treatment, and is transcribed into a non-coding neuronal RNA [20]. They found that the transcript was prominently increased in the small-type Kenyon cells of the dancers and foraging bee brains, suggesting involvement of the gene in foraging behavior. However, its involvement during the course of foraging had never been studied before. This is the first report on transient overexpression of *kakusei* during foraging. We have further examined the function of the gene before and after foraging. After food reward *kakusei* was immediately overexpressed over a short period of time from 15 to ca. 45 to 60 min, and then started decreasing. Interestingly, a short time overexpression at 15min and no further increase of the gene level without reward reveals the gene’s involvement in the learned food reward. This indicates that food reward is essential for the increase and sustaining of higher *kakusei* levels during foraging, thus underlying the role of *kakusei* in associative learning during foraging. This finding of *kakusei* involvement in learning and memory is further supported by the earlier report by Kiya et al., 2007 [20] as they reported overrepresentation of *kakusei* among the re-orienting Kenyon cells of bees assumed to be foragers. This is in line with the notion that re-orientation flight was the mechanism of flight behavior that was practiced by those bees for memorizing the hive location, thus indicating a role of *kakusei* in the information integration during foraging flight, which is an essential part for dance communication [20]. We suppose that the *kakusei* gene might also be involved in social interaction and communication among the foraging bees, linking through different molecular pathways as these behaviors were clearly observed among the foragers over the two hours of foraging. Our recent study [12] identified *egr-1*and *hr38*, and downstream genes such as *ecr, dopecr, ddc*, and *dopr2* that are involved in the ecdysteroid pathway and dopaminergic pathway respectively. Further, possible coordination of *kakusei* and the two IEGs *egr-1* and *hr38* during foraging behavior is quite possible and thereby to their downstream molecular pathways that include dopamine and ecdysteroid pathway and also in learning and memory as all the three genes demonstrated associative learning during foraging. Since *kakusei* is a non coding gene, this study also clearly support the dynamic role of non coding genes in daily behavior.

## Conclusion

Previous reports have already indicated involvement of immediate early genes could be used as neural marker and their association in social behaviors such as learning and memory or memory processing. Our recent paper by Singh et al., 2018 [12], has further supported the evidence as the two IEGs *egr-1* and *hr38* immediately and transiently expressed during honey bee foraging and further their involvement in memory processing or associative learning. The present finding on recently discovered IEG *kakusei* is an addition into the circle, thus increases the number of IEGs and ultimately more IEG choices available that could be used as search tool in finding molecular and cellular signaling underlying social behaviors, using honey bee foraging as model system. How the three IEGs *egr-1, hr38* and *kakusei* coordinate to complete the foraging task via their specific role in the learning, memory and their roles in communication and interaction and the underlying molecular and cellular pathways that link to each of this specific behavior will provide a thorough understanding of these behaviors at molecular and cellular levels. Thereby the hidden mechanisms of social behavior in honey bees could be opened up and mapped which could be further translated to higher animals and man [28].

## Acknowledgement

Dr. Axel Brockmann is gratefully acknowledged for providing the facilities in his lab to work and complete this project. Dr. Brockmann also read the manuscript and provided valuable comments. Ms. Neha Tanwar and Dr. Sophia Liyang kindly supported during experimental sample preparation. The Council of Scientific and Industrial Research (CSIR), Govt. of India and National Centre for Biological Sciences (NCBS), TIFR, India, are gratefully acknowledged for the fellowships provided to Dr. Asem Surindro Singh for successfully completing this project.

## Author’s Contribution

Dr. Asem Surindro Singh did the conceptualization of hypothesis, designing and conducting experiments, data analysis as well as manuscript writing. Dr. Pamela Cappelletti and Dr. Marco Feligioni contributed in manuscript editing.

